# Identification of SASP-associated biomarkers and regulatory mechanisms in diabetic foot ulcers based on transcriptomics and experimental validation

**DOI:** 10.64898/2025.12.09.693305

**Authors:** Tao Liu, Zhijun He, Yan Li, Xiaotao Wei, Xingzhang Yao, Wen Chen, Jinpeng Li, Fei Li, Bihui Bai

## Abstract

**Background:** Diabetic foot ulcers (DFU) constitute a major complication arising from diabetes mellitus. Emerging research findings have underscored the pivotal contribution of cellular senescence to the pathophysiological development of wounds exhibiting delayed healing; nevertheless, the molecular regulatory networks governing this process in DFU pathogenesis remain incompletely elucidated

**Methods:** DFU-related transcriptomic datasets were acquired from public databases. Biomarkers were identified through machine learning algorithms combined with expression validation. To evaluate the diagnostic predictive capacity of identified biomarkers for DFU, a nomogram prediction model was developed and validated. Further analyses included Gene Set Enrichment Analysis (GSEA), molecular network regulation, immune infiltration profiling, and drug prediction. Subsequently, the expression profiles of these biomarkers were quantitatively assessed through reverse transcription quantitative PCR (RT-qPCR) methodology.

**Results:** Four feature genes were identified using machine learning. Among them, CXCR2 and SIGLEC7 passed expression validation and were significantly up-regulated in DFU groups across two datasets (*p* < 0.05), confirming their potential as biomarkers. A nomogram constructed based on these two biomarkers demonstrated high discriminative accuracy and good calibration (decision curve net benefit > 0, AUC = 0.99). GSEA indicated that CXCR2 and SIGLEC7 were both enriched in the cytokine-cytokine receptor interaction pathway as well as other functionally associated pathways. Molecular regulatory network analysis indicated that both CXCR2 and SIGLEC7 were predicted to be regulated by transcription factors (TFs) such as SOX2. Immune infiltration analysis showed a significant strong positive correlation between CXCR2 and activated mast cells (correlation coefficient (cor) = 0.72, *p* < 0.001), and between SIGLEC7 and macrophages M0 (cor = 0.71, *p* < 0.001). Drug prediction identified 12 drugs targeting CXCR2 (e.g., Indoprofen) and one drug targeting SIGLEC7 (N-acetyl-alpha-D-glucosamine). Validation through Reverse Transcription quantitative Polymerase Chain Reaction (RT-qPCR) revealed that CXCR2 and SIGLEC7 expression levels were markedly increased in the DFU group, showing statistically significant differences (*p*<0.05), which aligned with the bioinformatics analysis outcomes

**Conclusion:** This study identified CXCR2 and SIGLEC7 as SASPRG-related biomarkers in DFU through bioinformatic analysis, providing new theoretical support and a basis for early diagnosis and treatment of DFU.

## 1. Introduction

Diabetes mellitus (DM) is a global health challenge, and diabetic foot ulcer (DFU) stands out as a severe and prevalent complication among its associated foot diseases(1). DFU, defined as skin breakdown involving at least the epidermis and part of the dermis below the ankle in diabetic patients, often coexists with peripheral neuropathy and/or peripheral arterial disease (2). The characteristic clinical manifestations include ulceration on the sole (commonly in the forefoot weight-bearing area), blood blisters/ulcers under calluses, exudation, and foul odor. Neuropathy leads to painless progression due to reduced pain sensation, while ischemia is manifested as cold feet, weak pulses, and poor-quality granulation tissue (3–4). Epidemiologically, DFU affects approximately 18.6 million people worldwide annually, serving as a precursor to most lower-limb amputations. The condition presents a serious risk to patients’ lives and quality of life, with a 5-year mortality rate of approximately 30% and over 70% in major amputation cases, along with high recurrence rates of 42% within a year and 65% within five years post-healing (5).The current approach to treating DFU involves a multidisciplinary strategy, focusing on surgical debridement, off-loading, addressing lower-limb ischemia and foot infections, and early referral for comprehensive care (6). However, due to the complex etiology and the often irreversible nature of blood supply and nerve damage, early screening and individualized interventions remain limited, resulting in low healing rates, high recurrence rates, and suboptimal long-term prognoses (7–8). Therefore, uncovering the underlying molecular mechanisms and identifying sensitive and specific biomarkers are crucial for improving the early detection and optimizing clinical intervention strategies for DFU.

Cellular senescence, marked by a largely irreversible halt in cell division due to different internal and external stress factors while preserving metabolic function, leads to the development of the senescence-associated secretory phenotype (SASP)(9). Containing pro-inflammatory cytokines, chemokines, growth factors, and proteases, SASP serves two roles: it helps with tissue repair in healthy conditions but contributes to chronic inflammation and tissue damage in pathological situations (10–11). Increasing evidence has linked SASP to the development and progression of DFU. In the microenvironment of DFU, factors such as high glucose, oxidative stress, and local ischemia-infection induce the accumulation of senescent keratinocytes, fibroblasts, and endothelial cells, which continuously secrete SASP, leading to persistent inflammation, inhibited matrix repair, re-epithelialization, and angiogenesis, thus keeping the wound in a non-healing state (12–14). Transcriptomic analysis of the edges of human DFU ulcers has shown up-regulation of multiple senescence markers (e.g., CDKN1A, IL1A, CXCL8, MMP10, SERPINE1) compared to non-ulcerated diabetic foot skin, indicating the involvement of the senescence-SASP network in the maintenance of DFU (15). Nevertheless, the specific regulatory mechanisms of SASP in DFU remain unclear and require further in-depth exploration.

This research sought to fill these knowledge gaps by systematically combining transcriptomic data from various public databases with weighted gene co-expression network analysis (WGCNA) and machine learning methods. By doing so, we aimed to identify potential biomarkers, construct nomograms, analyze molecular network regulation and immune infiltration, with a particular focus on the role of SASP in DFU. Our findings are expected to provide new insights into the early diagnosis and clinical management of DFU, ultimately improving patient outcomes.

## 2. Materials and methods

### 2.1 Data source

Two publicly available transcriptomic datasets for DFU were screened and downloaded from the Gene Expression Omnibus (GEO) database (https://www.ncbi.nlm.nih.govgeo/) for use as training and validation sets. The training set, GSE199939 (platform GPL199939), contained ten DFU skin tissue samples and 11 non-diabetic foot skin tissue samples. The validation set, GSE80178 (platform GPL16686), consisted of six DFU foot ulcer tissue samples and three non-diabetic foot ulcer tissue samples. Additionally, 125 SASP-related genes (SASPRGs) were obtained from the literature (16) **(Table S1)**.

### 2.2 Differential expression analysis

For the identification of differentially expressed genes (DEGs) comparing DFU specimens with control specimens within the training dataset GSE199939, differential expression profiling was conducted utilizing the “DESeq2” package (v 1.40.2; 17). The filtering thresholds were set as |log₂Fold Change (FC)| ≥ 2 and *p*.adj < 0.05. For visualization purposes depicting the global distribution patterns of differentially expressed genes, volcano plots alongside heatmaps were created through the “ggplot2” package (v 3.4.1; 18) and “ComplexHeatmap” package (v 2.16.0; 19), respectively. The leading ten genes exhibiting upregulation and downregulation, ordered according to |log2FC| values, were annotated within these visualizations.

### 2.3 Single-sample gene set enrichment analysis (ssgsea) for calculating SASPRGs scores

To examine the relationship between SASPRGs and DFU pathogenesis, the ssGSEA algorithm integrated within the “GSVA” package (v 1.50.0; 20) was utilized to compute SASPRGs scores across all specimens in the training dataset GSE199939, referencing the previously described 125 SASPRGs. Differences in SASPRGs scores between DFU and control samples were compared using the Wilcoxon test (*p* < 0.05), and a violin plot was generated using the “ggplot2” package (v 3.4.1) to visualize the results.

### 2.4 Weighted gene co-expression network analysis (WGCNA)

To identify SASPRGs module genes, WGCNA was performed on all samples in the training set GSE199939, using SASPRGs scores as traits to further screen for SASPRGs-related module genes. Co-expression network construction was achieved using the “WGCNA” package (v 1.71;21). The goodSampleGene function was utilized to screen for outlier specimens and evaluate the overall correlations among samples. The pickSoftThreshold function was implemented to establish the appropriate soft threshold (power) for sample clustering: an optimal power value ranging from 1 to 20 was chosen to set the threshold governing gene correlation; associations between the soft threshold and the scale-free network fit index R², along with associations between the soft threshold and mean connectivity, were constructed (utilizing R ² ≥ 0.80 as the selection criterion). Following the selection of the appropriate soft threshold, the adjacency matrix derived from sample clustering was converted into a Topological Overlap Matrix (TOM). Hierarchical clustering was then performed using the hierarchicalCluster function to identify modules. To further screen module genes strongly associated with SASPRGs scores, correlation and p-value analysis methods from the “WGCNA” package (v 1.71) were applied. Modules demonstrating significant correlation with SASPRGs scores (|correlation coefficient (cor)| > 0.3, *p* < 0.05) were recognized, with the module exhibiting the maximum absolute correlation coefficient coupled with the minimum *p*-value being designated as the pivotal module. The geneModuleMembership and geneTraitSignificance functions were used to analyze Module Membership (MM) and Gene Significance (GS) for module genes. Genes meeting |MM| > 0.8 and |GS| > 0.2 were selected as key module genes (22), denoted as SASPRGs module genes (MSASPRGs), for subsequent analysis.

### 2.5 Identification of Candidate Genes (CGs)

To further screen for differentially expressed CGs associated with SASPRGs, an intersection analysis was conducted between the previously filtered DEGs and MSASPRGs. A Venn diagram was plotted using the “ggvenn” package (v 0.1.10; 23) to visualize the overlap between the two gene sets. The overlapping genes were extracted and defined as CGs.

### 2.6 Enrichment analysis and protein-protein interaction (PPI) analysis

The Gene Ontology (GO) provides a systematic description of gene functional attributes. TThe Kyoto Encyclopedia of Genes and Genomes (KEGG) serves as an extensively utilized repository for investigating metabolic and signaling pathway dynamics, facilitating comprehension of gene functionality across intricate biological systems. To gain deeper insights into the potential biological functions and pathways through which CGs contribute to DFU development and progression, GO and KEGG enrichment analyses were conducted utilizing the “clusterProfiler” package (v 4.2.2; 24), establishing a significance cutoff at p.adjust < 0.05. The leading ten significantly enriched GO terms alongside KEGG pathways were graphically represented via the “ggplot2” package (v 3.4.1) according to p-values, thereby illuminating underlying biological mechanisms. To examine protein-level interactive relationships among CGs, a PPI network was assembled through STRING (https://string-db.org/, v 12.0) employing a confidence cutoff > 0.4. Visualization of the generated network was achieved utilizing Cytoscape software (v 3.8.2; 25).

### 2.7 Machine learning-based screening of feature genes

For subsequent identification of DFU-associated biomarkers from CGs, two machine learning methodologies were systematically employed for feature selection utilizing the training set GSE199939. Initially, the Least Absolute Shrinkage and Selection Operator (LASSO) algorithm incorporating 5-fold cross-validation was executed via the “glmnet” package (v 4.1.8; 26). Configuration parameters specified family = “binomial”, with the minimal lambda value being chosen as the optimal cutoff for executing LASSO logistic regression, culminating in the determination of LASSO feature genes. Second, the XGBoost algorithm was employed using the “xgboost” package (v 2.0.3.1; 27) for feature selection of CGs. Samples were divided into “control” and “disease” groups, and a binary classification model with the objective function “binary:logistic” was constructed. The number of boosting rounds (nround) was set to 25, maximum tree depth (max-depth) to two, and learning rate (eta) to 0.3. After training, the xgb.importance function was used to evaluate the importance (Gain value) of each gene, and a feature importance bar plot was generated. Finally, XGBoost feature genes were screened based on this evaluation. To enhance the reliability of feature gene selection, the results from both algorithms were intersected using the “ggvenn” package (v 0.1.10). The overlapping genes were defined as final feature genes for subsequent analysis.

### 2.8 Expression validation for biomarker screening

To comprehensively verify the transcriptional variations of candidate genes distinguishing DFU patients from healthy controls and to characterize biomarkers demonstrating robust expression profiles, differential gene expression analyses were conducted employing the non-parametric Wilcoxon rank-sum test across both the discovery cohort GSE199939 and the independent validation cohort GSE80178. Genes exhibiting concordant directional changes and achieving statistical significance (*p*-value threshold < 0.05) in both cohorts were retained for subsequent analyses. These consistently dysregulated genes were designated as the diagnostic biomarkers for the present investigation. Visualization of expression patterns across experimental groups was accomplished utilizing the “ggplot2” visualization framework (version 3.4.1), enabling intuitive representation of expression heterogeneity between conditions.

### 2.9 Construction and evaluation of the nomogram

To comprehensively assess the discriminatory capacity of identified biomarkers in DFU diagnosis, a predictive nomogram was developed leveraging the “rms” statistical package (version 6.8.1; 28). This nomogram integrated the aforementioned biomarker panel with the complete sample population from the discovery dataset GSE199939 to generate individualized DFU risk probability estimates. The nomogram assigns a score to each biomarker, and the sum of these scores corresponds to the total points. The DFU incidence is then inferred based on the total points, with higher total points indicating a higher probability of DFU. The calibration curve of the nomogram was generated using the “rms” package (v 6.8.1) to assess the accuracy of the predictions. Model calibration was evaluated through calibration plot analysis, wherein proximity of the calibration curve to the 45-degree reference line indicates superior concordance between predicted probabilities and actual outcomes. The Hosmer-Lemeshow goodness-of-fit test (HL test) served as the primary metric for assessing model calibration, quantifying discordance between predicted and empirically observed values. An HL test *p*-value exceeding 0.05 signifies adequate model calibration, indicating the absence of statistically significant deviation between model predictions and observed outcomes, thereby confirming satisfactory model performance. Furthermore, decision curve analysis (DCA) was performed using the “ggDCA” package (v 1.1; 29), and a DCA curve was plotted. A net benefit value > 0 indicates effective predictive performance of the model. Subsequently, the “pROC” package (v 1.18.5; 30) was used to generate ROC curves. The AUC value was calculated to evaluate the predictive performance of the nomogram model for disease samples in the training set, with an AUC > 0.7 considered indicative of good predictive value.

### 2.10 Chromosomal localization and gene set enrichment analysis (GSEA)

Chromosomal localization analysis of the identified biomarkers was performed utilizing the “OmicCircos” visualization package (version 1.38.0;31), which enabled precise mapping of these molecular signatures to their respective genomic coordinates across human chromosomes. To elucidate the underlying regulatory networks and biological processes implicated by these diagnostic markers, GSEA targeting KEGG pathways was implemented through integration of multiple bioinformatics tools, including the “clusterProfiler” analytical framework (v 4.2.2), the “enrichplot” visualization suite (v 1.18.3; 32), and the “msigdbr” database interface package (v 7.5.1; 33). This enrichment analysis was conducted on the complete cohort of DFU patients and healthy controls from the discovery dataset GSE199939. Reference gene set collections for Homo sapiens were retrieved from the Molecular Signatures Database (MSigDB, https://www.gsea-msigdb.org/gsea/msigdb), specifically leveraging the curated “CP:KEGG” pathway compendium within the C2 canonical pathways category, thereby facilitating systematic identification of significantly enriched biological pathways associated with the biomarker panel. Pairwise correlation analyses between each biomarker and the entire transcriptome were quantified through Spearman’s rank correlation methodology implemented via the “psych” statistical package (v 2.1.6; 34). Subsequently, all genes were hierarchically arranged in descending order according to their correlation coefficient magnitudes, generating a comprehensively ranked gene list for downstream enrichment interpretation. The top ten pathways were selected and displayed according to p-values. Significantly enriched pathways for each biomarker were identified using the following criteria: |normalized enrichment score (NES)| > 1, *p* < 0.05, and false discovery rate (FDR) < 0.25. These pathways were used to infer potential molecular mechanisms and pathological processes involved.

### 2.11 Molecular network regulation

To comprehensively elucidate the molecular regulatory networks governing DFU pathogenesis and disease progression, upstream transcriptional regulators controlling the expression of identified biomarkers were systematically inferred through the ChEA3 computational platform (https://maayanlab.cloud/chea3/) integrated within the NetworkAnalyst bioinformatics suite. This transcription factor (TF) prediction analysis was complemented by post-transcriptional regulatory element identification, wherein the get-multimir algorithmic function embedded in the “multiMiR” R package (version 1.16.0; 35) was deployed to computationally predict microRNA (miRNA) species demonstrating putative targeting interactions with the biomarker transcripts. Finally, a TF-mRNA-miRNA regulatory network was constructed using Cytoscape software (v 3.8.2).

### 2.12 Immune infiltration analysis

Characterization of immune microenvironment heterogeneity distinguishing DFU lesions from normal tissue was accomplished through comprehensive immune cell composition profiling. Specifically, the CIBERSORT deconvolution algorithm (36) was applied to the complete specimen cohort from the discovery dataset GSE199939, encompassing both DFU patients and healthy controls, to quantitatively estimate the relative abundance fractions of twenty-two distinct immune cell subpopulations within each individual sample. The compositional landscape of these immune cell subtypes was graphically represented utilizing the “ggplot2” graphics framework (v 3.4.1), enabling intuitive visualization of infiltration magnitudes across the immune cell repertoire. Statistical comparison between DFU and control groups was conducted via the non-parametric Wilcoxon rank-sum test within the discovery cohort to pinpoint immune cell populations exhibiting statistically significant alterations, employing a stringent significance cutoff of p-value < 0.05. Distribution patterns of significantly altered immune populations were illustrated through boxplot representations generated via the “ggplot2” package (v 3.4.1). To systematically dissect the interrelationships among differentially abundant immune cell populations, as well as to elucidate potential associations linking biomarker expression profiles with immune infiltration patterns, pairwise Spearman’s rank-order correlation matrices were computed across the entire sample set from the training cohort GSE199939 using the “psych” statistical computing package (v 2.1.6). Correlations with *p* < 0.05 and |cor| > 0.3 were considered significant. The resultant correlation architecture was visually rendered through heatmap representations constructed with the “corrplot” visualization package (version 0.92;37).

### 2.13 Drug prediction and molecular docking

To explore potential drugs related to DFU, the biomarkers were imported into the DrugBank database (https://go.drugbank.com/) to identify existing drugs or small organic molecules capable of targeting the biomarkers. The resulting compounds were designated as candidate drugs. To further validate the interactions between candidate drugs and biomarkers, molecular docking was performed to predict binding affinity and mechanisms of action, enabling the screening of more suitable drugs. Chemical structural information for candidate pharmaceutical compounds was systematically retrieved from the publicly accessible PubChem repository (https://pubchem.ncbi.nlm.nih.gov/), whereas crystallographic three-dimensional conformations of protein products corresponding to the identified biomarkers were acquired from the Research Collaboratory for Structural Bioinformatics Protein Data Bank (PDB, https://www.rcsb.org/). Protein preparation, including dehydration, phosphate removal, and dehydrogenation, was conducted using PyMOL (https://pymol.org/2/; v 3.1.3; 38). Finally, molecular docking was carried out using AutoDock Vina (v 1.5.7;39). A docking score ≤ −5 kcal/mol indicated strong binding affinity. This analysis provided structural insights into the mechanisms of action of the potential drugs.

### 2.14 Reverse transcription quantitative polymerase chain reaction, (RT-qPCR)

To investigate the differences in the expression of biomarkers between DFU and control samples, this study collected blood samples from DFU patients (n = 6) and normal controls (n = 6). Healthy control skin samples were obtained from the lower limb area during internal fixation removal surgery. Tissue samples from patients with DFU were collected from the wound edges during debridement. Total ribonucleic acid was isolated from individual peripheral blood specimens utilizing the Trizol reagent-based extraction protocol (Vazyme, catalog number R401-01, China). Subsequently, the purified RNA templates underwent reverse transcription into first-strand complementary deoxyribonucleic acid (cDNA) employing the HP All-in-one qRT Master Mix II RT203-Ver.1 for qPCR with integrated genomic DNA elimination functionality (gDNA digester plus) (YoungGen, lot number 24Y0124, China). Quantitative real-time polymerase chain reaction (RT-qPCR) amplification was executed utilizing the 2 × Universal Blue SYBR Green qPCR Master Mix system (Servicebio, product code G3326-05, China). Statistical evaluation of differential expression patterns between experimental cohorts was accomplished through Student’s t-test methodology with a predetermined significance threshold of *p* < 0.05. Relative quantification of biomarker transcript abundance within each clinical specimen was computed via the comparative 2^-ΔΔCt^ algorithm. Oligonucleotide primer sequences specific for biomarker amplification were commercially synthesized by Sangon Biotech Co., Ltd. (Shanghai, China) with detailed sequences provided in Table 2. Experimental reproducibility was ensured through triplicate technical replication of each biological specimen. Ethical compliance for this investigation was secured through formal approval granted by the Institutional Review Board of Gansu Provincial Hospital of Traditional Chinese Medicine (approval identifier: GSZY-2025-044-01). Glyceraldehyde-3-phosphate dehydrogenase (GAPDH) served as the endogenous housekeeping reference gene for normalization purposes. Comprehensive statistical evaluation and graphical visualization of experimental datasets were accomplished using the GraphPad Prism software platform (v 10.1.2) (40).

### 2.15 Statistical analysis

Computational bioinformatic procedures were implemented within the R statistical computing environment (v 4.2.2). Between-group comparative analyses were performed employing both the non-parametric Wilcoxon rank-sum test and the parametric Student’s t-test, with statistical significance uniformly defined at a probability threshold of *p* < 0.05; specifically, p-values falling below this 0.05 cutoff were interpreted as indicating statistically meaningful differences.

## 3. Results

### 3.1 Identification of DEGs

In the differential expression analysis, expression differences between DFU samples and control samples in the training set were compared, with screening criteria set as |log₂FC| ≥2 and adj *p* < 0.05. Through differential expression analysis, 1,806 differentially expressed genes were identified from the dataset, comprising 537 genes showing elevated expression levels and 1,269 genes exhibiting reduced expression patterns (**Table S3**). Volcano plots and heatmap representations were constructed to illustrate the spatial distribution patterns and statistical significance of these differentially expressed genes, with particular emphasis on highlighting the ten genes demonstrating the most pronounced upregulation and downregulation profiles.nd heat maps, with the top ten most significantly up-regulated and down-regulated genes labeled. Compared to control samples, TDO2, IL11, ERFE, LINC01614, CXCL13, MMP3, WTAPP1, MMP1, CXCL5, and MMP13 were significantly highly expressed in DFU samples, while KRT1, KRT14, KRT77, CDHR1, KRT5, MAL2, SFN, DSC3, EVPL, and TNS4 were significantly down-regulated **(Fig1A-B)**.

**Figure 1.** Identification of differentially expressed genes (A) Volcano plot of differential expression analysis between the DFU group and the control group. (B) Heatmap of the top 10 up-regulated and down-regulated genes in transcriptomic differential analysis between DFU samples and control samples.

### 3.2 Identification of MSASPRGs

Based on the ssGSEA scores of SASPRGs in DFU, significant differences were observed between DFU and control samples (*p* < 0.001) **(Fig 2A)**. Hierarchical clustering methodology was subsequently applied to the training cohort samples through WGCNA to evaluate inter-sample correlations across the entire dataset. The results showed no significant outlier samples **(Fig 2B)**. Upon establishing the coefficient of R^2^ at 0.8, the analysis revealed that a soft-thresholding power of 21 represented the optimal parameter value, thereby satisfying the criteria for scale-free topology network construction (**Fig 2C**). Following hierarchical clustering and correlation screening between modules and SASPRGs scores, eight key modules were ultimately identified (excluding the grey module) **(Fig 2D)**. To establish the association between SASPRGs and the modules, the module with the highest positive correlation-the green module (cor = 0.81, *p* = 8e-06)-was selected as the key module, containing 492 genes **(Fig 2E, Table S4)**. By applying thresholds of |MM| > 0.8 and |GS| > 0.2, 219 genes were screened as key module genes, designated as MSASPRGs **(Fig 2F, Table S5)**. This approach enabled the identification of critical functional gene groups from a large pool, revealing core biological processes.

**Figure 2.** Identification of MSASPRGs in DFU (A) Differences in SASPRGs scores between DFU samples and control samples. (B) Clustering dendrogram of all samples in the GSE199939 database. (C) Soft threshold screening: determination of the optimal soft threshold (left panel); network connectivity under different soft thresholds (right panel). (D) Module clustering dendrogram. The vertical axis represents the dissimilarity between genes or modules-lower height indicates higher similarity in expression patterns. The horizontal axis arranges all genes or modules based on clustering algorithms (e.g., hierarchical clustering). The color bar (typically blocks below) assigns each color to a co-expression module, with genes of the same color sharing similar expression patterns. (E) Heatmap of correlations between different key modules from WGCNA and SASPRGs. (F) Scatter plot of GS and MM for the key module.

### 3.3 Identification and functional analysis of CGs

The intersection of 1,806 DEGs obtained from differential expression analysis and 219 MSASPRGs yielded 25 CGs **(Fig 3A, Table S6)**. Functional enrichment analysis of the 25 CGs using GO terms revealed significant enrichment in 81 pathways (p < 0.05) **(Table S7)**. Major enriched biological processes included: myeloid leukocyte activation, leukocyte migration, leukocyte chemotaxis, myeloid leukocyte migration, granulocyte chemotaxis, leukocyte activation involved in immune response, cell activation involved in immune response, cell chemotaxis, granulocyte migration, and Fc receptor signaling pathway **(Fig 3B)**. KEGG pathway analysis identified 53 significantly enriched pathways (*p* < 0.05) **(Table S8)**, including neutrophil extracellular trap formation, viral protein interaction with cytokines and cytokine receptors, leishmaniasis, cytokine-cytokine receptor interaction, systemic lupus erythematosus, osteoclast differentiation, malaria, tuberculosis, legionellosis, and chemokine signaling pathway **(Fig 3C)**. These results indicated that the CGs were involved in biological processes such as immune and inflammatory responses, as well as cell differentiation and pathological remodeling. PPI analysis among the CGs was further performed, with nodes filtered at a confidence score > 0.4. The results indicated that CXCR2 interacts with multiple proteins, such as SELL, CXCL8, and CCR1 **(Fig 3D)**, revealing complex interaction relationships among the CGs and providing a structural basis for further investigation into disease regulatory mechanisms.

**Figure 3.** Identification and functional enrichment analysis of candidate genes (A) Venn diagram of MSASPRGs and DEGs. (B) GO enrichment analysis of candidate genes. (C) KEGG enrichment analysis of candidate genes. (D) PPI network of candidate genes.

### 3.4 Identification of biomarkers

Subsequently, to further identify feature genes, two machine learning methods were employed for gene screening. First, LASSO regression was used for variable compression and selection. At log(λ _min_) = −5.3215, the model’s prediction error (i.e., cross-validation error) reached its minimum, and nine LASSO feature genes were screened: CXCL8, CCR1, CXCR2, TNFRSF10C, MCEMP1, TREM1, SIGLEC7, FCGR1CP, and TREML2 **(Fig 4A-B)**. Second, six XGBoost feature genes were obtained using XGBoost **(Fig 4C)**, including FCGR3A, MCEMP1, SIGLEC7, CCR1, and CXCR2. Finally, by integrating the results of both methods and taking their intersection, four common feature genes were identified: CCR1, CXCR2, MCEMP1, and SIGLEC7 **(Fig 4D)**. These were defined as feature genes and served as core molecules for subsequent diagnostic and mechanistic studies.

**Figure 4.** Identification of biomarkers (A) Cross-validation curve of LASSO regression analysis. (B) LASSO coefficient profile plot. (C) Gene importance scores screened by XGBoost. (D) Venn diagram of core genes identified by LASSO and XGBoost machine learning algorithms. (E) Boxplot of candidate biomarker expression in the training set GSE199939. (F) Boxplot of candidate biomarker expression in the validation set GSE80178. \*\*\**p* < 0.001, \*\**p* < 0.01, \**p* < 0.05, ns: not significant.

To determine whether the expression patterns of these four feature genes were consistent across the training set GSE199939 and the validation set GSE80178, expression comparison analysis was performed to evaluate their differences between DFU samples and control samples. The results showed that CXCR2 and SIGLEC7 exhibited significant differences (*p* < 0.05) between the DFU and control groups in both datasets. Specifically, both CXCR2 and SIGLEC7 were significantly up-regulated in the DFU group **(Fig 4E-F)**, confirming their potential as biomarkers for subsequent research.

### 3.5 Nomogram prediction model based on CXCR2 and SIGLEC7

The ability of the biomarkers to predict DFU was evaluated. The results showed that an individual with a total score of 164 had a predicted DFU risk of 91.7% **(Fig 5A)**. The calibration curve analysis was conducted to verify the model’s forecasting capability, revealing substantial concordance between the nomogram-derived probability estimates and the ideal reference diagonal. Statistical assessment yielded a p-value of 0.924, markedly exceeding the 0.05 threshold, thereby demonstrating negligible discrepancy between projected and observed outcomes, which substantiated the nomogram’s robust predictive precision (**Fig 5B**). Additionally, DCA analysis illustrated that the model’s decision curve consistently maintained positive values throughout the threshold probability range, signifying favorable clinical net benefit. The model’s DCA curve was also positioned above the “All” and “None” lines, suggesting favorable predictive performance **(Fig 5C)**. Subsequently, ROC curve analysis showed an AUC value of 0.99, far exceeding the threshold of 0.7, further demonstrating excellent predictive capability **(Fig 5D)**. In summary, these findings confirm that the nomogram model is reliable and suitable for clinical prediction, providing a robust tool for the early diagnosis of DFU.

**Figure 5.** Construction and evaluation of the biomarker-based nomogram (A) Nomogram constructed using biomarkers. (B) Calibration curve. (C) Decision curve analysis (DCA). (D) ROC analysis of the model.

### 3.6 Chromosomal localization and GSEA analysis of CXCR2 and SIGLEC7

The specific chromosomal locations of CXCR2 and SIGLEC7 were explored. CXCR2 was found to be located on chromosome two, and SIGLEC7 on chromosome 19 **(Fig 6A)**. Subsequently, the potential biological functions of CXCR2 and SIGLEC7 in the pathogenesis of DFU were investigated using GSEA. Pathway enrichment analysis revealed that CXCR2 exhibited statistically significant enrichment across 75 distinct biological pathways, with all associations meeting the significance criterion (*p* < 0.05). The top ten pathways ranked by enrichment p-value included: Nod-like receptor signaling pathway, Cytokine-cytokine receptor interaction, Proteasome, Chemokine signaling pathway, Jak-Stat signaling pathway, Toll-like receptor signaling pathway, Natural killer cell mediated cytotoxicity, and Cell cycle **(Fig 6B, Table S9)**. SIGLEC7 was significantly enriched in 84 pathways (p < 0.05). The top ten pathways ranked by enrichment p-value included Proteasome, Cytokine-Cytokine Receptor Interaction, and Chemokine Signaling Pathway, among others (**Fig 6C, Table S10**).

**Figure 6.** Chromosomal localization and GSEA analysis of biomarkers (A) Chromosomal localization of biomarkers. (B) GSEA results for CXCR2. (C) GSEA results for SIGLEC7.

Both genes were co-enriched in pathways such as Chemokine-signaling-pathway, Leukocyte-transendothelial-migration,FC-gamma-R-mediated-phagocytosis, and Cytokine-cytokine-receptor-interaction, suggesting that CXCR2 and SIGLEC7 primarily influence the occurrence and development of DFU through inflammatory pathways.

### 3.7 Molecular network regulation and immune infiltration analysis of CXCR2 and SIGLEC7

TFs and miRNAs targeting the regulation of CXCR2 and SIGLEC7 were explored, and a TF-mRNA-miRNA network was constructed. The results revealed that SOX2, GATA2, TP63, and GATA3 were commonly predicted to regulate both CXCR2 and SIGLEC7, indicating that these two genes are co-regulated by shared upstream factors and thereby influence the development of DFU. Furthermore, 15 miRNAs were predicted to target CXCR2, including hsa-miR-593-5p, hsa-miR-224-5p, and hsa-miR-302d-3p. Four miRNAs were predicted to target SIGLEC7: hsa-miR-936, hsa-miR-30e-3p, hsa-miR-30d-3p, and hsa-miR-30a-3p**(Fig 7A)**. These findings suggest that CXCR2 and SIGLEC7 are involved in a complex TF-mRNA-miRNA regulatory network that modulates the progression of DFU. To gain deeper insights into the immune modulation pathways potentially orchestrated by the identified biomarkers (CXCR2 and SIGLEC7), comparative analysis demonstrated statistically significant disparities (*p* < 0.05) across eight distinct immune cell populations when contrasting the DFU cohort against control subjects. Notably, the DFU group exhibited markedly elevated infiltration levels of macrophages M0, whereas the abundance of resting dendritic cells and resting mast cells displayed pronounced reduction patterns (**Fig 7B, C**). Correlation analysis among the differentially infiltrated immune cells revealed that resting mast cells showed significant strong positive correlations with memory B cells (cor = 0.76, *p* < 0.05), activated NK cells (cor = 0.71, *p* < 0.05), and resting dendritic cells (cor = 0.63, *p* < 0.05). Resting dendritic cells exhibited significant strong negative correlations with resting NK cells (cor = −0.70, *p* < 0.05) and macrophages M0 (cor = −0.80, *p* < 0.05) **(Fig 7D)**. Further investigation into the correlation between the biomarkers and differentially infiltrated immune cells indicated that CXCR2 was significantly strongly positively correlated with activated mast cells (cor = 0.72, *p* < 0.001), while SIGLEC7 was significantly strongly positively correlated with macrophages M0 (cor = 0.71, *p* < 0.001) **(Fig 7E, Table S11)**. Overall, both CXCR2 and SIGLEC7 are closely associated with the immune microenvironment, suggesting their potential involvement in immune regulatory mechanisms and highlighting their significance for research on immune-related diseases and the immune microenvironment.

**Figure 7.** Network regulation and immune infiltration of biomarkers (A) TF-mRNA-miRNA regulatory network of biomarkers. (B) Heatmap of infiltration scores for 22 immune cell types in DFU and control groups. (C) Boxplot of immune cell infiltration for 22 immune cell types in DFU and control groups. (D) Heatmap of correlations among differentially infiltrated immune cells. (E) Heatmap of correlation analysis between biomarkers and differentially infiltrated immune cells.

### 3.8 Drug prediction and molecular docking analysis of CXCR2 and SIGLEC7

Existing drugs or small organic compounds capable of targeting CXCR2 and SIGLEC7 were explored. The results showed that 12 drugs were retrieved for CXCR2, including Indoprofen, Ibuprofen, and Clotrimazole, while one drug, N-acetyl-alpha-D-glucosamine, was identified for SIGLEC7 **(Fig 8A)**. Further investigation of the binding affinity between the biomarkers (CXCR2 and SIGLEC7) and the aforementioned small molecule drugs was conducted using molecular docking technology (docking score ≤ −5 kcal/mol). The results indicated that CXCR2 exhibited the strongest binding affinity with Clotrimazole, with a binding energy of −8.2 kcal/mol, and a binding energy of −6.8 kcal/mol with Ibuprofen. However, molecular docking diagrams revealed that Ibuprofen formed multiple hydrogen bond interactions with residues of CXCR2. SIGLEC7 showed a binding energy of −5.6 kcal/mol with N-acetyl-alpha-D-glucosamine, also forming multiple hydrogen bond interactions **(Fig 8B-D, Table 1)**. These findings collectively suggest that Clotrimazole, Ibuprofen, and N-acetyl-alpha-D-glucosamine may serve as potential therapeutic agents for DFU treatment.

**Figure 8.** Drug Prediction and Molecular Docking of Biomarkers (A) Drug-biomarker interaction network. (B) Molecular docking diagram of Clotrimazole with CXCR2. (C) Molecular docking diagram of Ibuprofen with CXCR2. (D) Molecular docking diagram of N-acetyl-alpha-D-glucosamine with SIGLEC7. Pink represents the protein structure of biomarkers, green represents small molecule drugs, and blue indicates amino acid residues involved in hydrogen bonding with the drugs.

### 3.9 Expression profiles of biomarkers

The results of RT-qPCR showed that in clinical samples, the expression levels of two biomarkers differed significantly between the DFU group and the control group (*p* < 0.05). Specifically, CXCR2 and SIGLEC7 were upregulated in the DFU group (Figure 9). The expression trends of these two genes in clinical samples were consistent with the results obtained from our bioinformatics analysis, indicating that the biomarkers identified through the analysis have high reliability.

## 4. Discussion

DFU is a severe complication of diabetes, closely linked to metabolic disorders, neuropathy, angiopathy, infection, and immune dysfunction induced by diabetes (41). Despite advances in clinical management, DFU treatment still faces limitations in improving healing rates and reducing recurrence (42).Growing evidence underscores the significant impact of cellular senescence in the development of chronic wounds, and targeting senescence or SASP appears promising for enhancing chronic wound healing. (43). To address this, our study analyzed transcriptomic data from the GEO database, integrating differential expression analysis, WGCNA, machine learning, and RT-qPCR validation. We identified two SASP-related diagnostic biomarkers, CXCR2 and SIGLEC7, constructed a nomogram to explore <u>the role of these two biomarkers</u> in DFU risk prediction, and analyzed their involved biological pathways and immune infiltration. This study offers fresh understanding of the molecular processes connecting SASP to DFU, aiding in early diagnosis and treatment.

This research identified two genes related to SASP, namely C-X-C motif chemokine receptor 2 (CXCR2) and sialic acid-binding Ig-like lectin 7 (SIGLEC7), as possible diagnostic markers for DFU using integrated bioinformatics and RT-qPCR validation, with both genes significantly upregulated in DFU tissues compared to controls (consistent between bioinformatic predictions and experimental results). CXCR2, a transmembrane G-protein-coupled receptor predominantly expressed on neutrophils, endothelial cells, and fibroblasts, binds ELR-motif-containing CXC chemokines (e.g., CXCL1, CXCL8/IL-8) to activate PLC-IP3/Ca²⁺, PI3K-AKT, and ERK/p38-MAPK pathways, regulating immune cell chemotaxis and inflammation (44–45). Notably, SASP is enriched with CXCR2 ligands (e.g., CXCL8), Creating an autocrine/paracrine loop that enhances NF-κB and p16/p21 signaling to reinforce cellular senescence—interrupting this loop has been suggested as a method to reduce chronic inflammation(46–47). In DFU, single-cell and spatial transcriptomics have revealed persistent activation of inflammatory chemotaxis networks, driven by neutrophils that promote matrix degradation and repair failure (48). Previous studies also highlighted a “senescence-CXCR2-inflammation” coupling in diabetic wounds: while transient CXCR2 activation facilitates early debridement, sustained overactivation exacerbates chronic inflammation (49–50). Our finding of CXCR2 upregulation in DFU supports a pathogenic model: hyperglycemia/ischemia/biofilms induce cellular senescence and SASP overproduction, which activates CXCR2 on neutrophils and endothelial cells—this triggers excessive neutrophil infiltration, neutrophil extracellular trap formation (NETosis), and MMP release, ultimately impairing re-epithelialization and angiogenesis via a “SASP-CXCR2-chronic inflammation” positive feedback loo<u>p, rendering DFUs difficult to heal over the long</u> <u>term.</u>

SIGLEC7 (also known as CD328), an inhibitory sialic acid-binding lectin on NK cells, macrophages, and monocytes, contains an ITIM motif that suppresses NK cell degranulation and cytotoxicity upon binding α2-8 disialylated ligands (e.g., ganglioside GD3) (51–52). Recent studies showed that senescent cells upregulate GD3 (via ST8SIA1) to form a “senescence immune checkpoint,” inhibiting NK cell-mediated senescent cell clearance and exacerbating fibrosis (53–54). DFU is characterized by accumulated senescent cells, and their clearance accelerates wound healing in diabetic mice (55). Our observation of SIGLEC7 upregulation in DFU suggests a mechanism: hyperglycemia/ischemia-induced SASP increases GD3 expression on local cells, enhancing SIGLEC7-mediated NK cell inhibition—this reduces senescent cell clearance, sustains SASP release, and traps wounds in a chronic inflammatory state. Notably, CXCR2 and SIGLEC7 have not been previously linked to DFU pathogenesis; our study is the first to validate their diagnostic potential and mechanistic roles, filling gaps in current understanding of SASP-driven DFU progression.

GSEA and KEGG analyses showed that CXCR2 and SIGLEC7 are jointly enriched in the chemokine signaling pathway, leukocyte transendothelial migration, FcγR-mediated phagocytosis, and cytokine-cytokine receptor interaction, which are crucial pathways in DFU pathogenesis.The chemokine signaling pathway, composed of CCL/CXCL chemokines and GPCRs (e.g., CXCR2), orchestrates immune cell trafficking and activation, with its precision regulating tissue immune homeostasis (56–57). In DFU, this pathway is dysregulated: hyperglycemia/ischemia stimulates keratinocytes and endothelial cells to secrete CXCL1/CXCL8, driving excessive neutrophil recruitment (58–59). For CXCR2, this overactivation exacerbates “inflammation-tissue destruction” feedback via neutrophil-derived ROS and proteases; for SIGLEC7, upregulation suppresses NK cell function, sustaining chemokine overproduction (e.g., CCL2) and monocyte/macrophage retention (60–61). Together, these disruptions lock wounds in the inflammatory phase, inhibiting ECM repair and angiogenesis (62).Leukocyte transendothelial migration mediates immune cell infiltration into wounded tissues. In DFU, hyperglycemia impairs endothelial barrier function, while CXCR2 upregulation enhances neutrophil-endothelial adhesion—this promotes abnormal leukocyte extravasation and tissue damage (63–64). Meanwhile, SIGLEC7 further dysregulates this process by inhibiting macrophage phagocytosis (via FcγR-mediated phagocytosis pathway), reducing pathogen clearance and resolving inflammation (64). Taken together, These pathway disruptions collectively contribute to DFU chronicity.

The analysis of immune infiltration revealed an increase in M0 macrophages and a decrease in resting dendritic cells (DCs) and mast cells in DFU, with significant correlations between biomarkers and immune cells: CXCR2 positively correlated with activated mast cells (r=0.72, *p*<0.001), and SIGLEC7 positively correlated with M0 macrophages (r=0.71, *p*<0.001). M0 macrophages are immature precursors that polarize to M1 (pro-inflammatory) or M2 (pro-repair) phenotypes—their accumulation in DFU indicates impaired polarization, as sustained M0 retention amplifies chronic inflammation and inhibits re-epithelialization (66–67). The positive correlation between SIGLEC7 and M0 macrophages suggests SIGLEC7 may suppress macrophage activation via ITIM signaling, further blocking M2 polarization (68). Resting DCs support immune balance through antigen presentation; their decrease in DFU hinders T cell activation and pathogen elimination, compromising adaptive immune responses (69–70). Resting mast cell depletion reflects excessive activation, as mast cell degranulation releases pro-inflammatory mediators (e.g., histamine) that exacerbate tissue damage and microcirculatory dysfunction (71–72). The correlation between CXCR2 and activated mast cells further supports a “CXCR2-mast cell” inflammatory loop: CXCR2 ligands (e.g., CXCL8) trigger mast cell degranulation, while mast cells secrete additional CXCR2 ligands, amplifying inflammation (73–74). Drug prediction identified 12 compounds targeting CXCR2 (e.g., indoprofen, ibuprofen, clotrimazole) and 1 targeting SIGLEC7 (N-acetyl-α-D-glucosamine, GlcNAc), with clinical relevance for DFU management. Clotrimazole, an imidazole antifungal, reduces DFU infection risk by clearing superficial fungi (e.g., tinea pedis), a common DFU trigger (75–76). Ibuprofen, delivered via foam dressings, alleviates DFU-associated pain and improves dressing tolerance—critical for patient compliance (77–78). GlcNAc, a component of chitosan dressings, enhances DFU healing via moisturization, antibacterial activity, and promotion of angiogenesis: a 2024 RCT (CHITOWOUND) showed chitosan gel significantly increased DFU area reduction (92% vs. 37% in controls, p = 0.008) (79). These agents provide actionable targets for DFU management, though further validation is needed.

This study has limitations: (1) Analyses relied on public GEO datasets, limiting sample size and generalizability; (2) Biomarker functions were not validated in cell/animal models; (3) Drug predictions lacked experimental verification. Future work will: (1) Validate CXCR2/SIGLEC7 roles in DFU using diabetic mouse models and primary human cells; (2) Test candidate compounds in preclinical studies; (3) Expand cohorts to confirm diagnostic utility.

In conclusion, our study identifies CXCR2 and SIGLEC7 as SASP-related diagnostic biomarkers for DFU, clarifies their mechanistic roles via pathway and immune infiltration analyses, and proposes candidate therapeutics—providing a foundation for improved DFU diagnosis and treatment.

## AUTHOR CONTRIBUTIONS

The contributions of all authors must be stated.Tao Liu: Data curation (equal); formal analysis (equal); investigation (equal); writing – original draft (equal).Zhijun He: Data curation (equal); formal analysis (equal); investigation (equal).Yan Li: Conceptualization (lead); supervision (lead); funding acquisition (lead); writing – review and editing (lead).Xiaotao Wei: Data curation (equal); project administration (equal).Xingzhang Yao: Data curation (equal); project administration (equal).Wen Chen: Data curation (equal); validation (equal).Jinpeng Li: Data curation (equal); validation (equal).Fei Li: Data curation (equal); formal analysis (equal).Bihui Bai: Data curation (equal); formal analysis (equal); writing – review and editing (equal).

## ACKNOWLEDGMENTS

The authors would like to express their gratitude to the staff of the Department of Foot and Ankle Orthopedics and Department of Orthopedic Repair and Reconstruction at Gansu Provincial Hospital of TCM for their support in sample collection and clinical data provision. The authors also thank the contributors of the Gene Expression Omnibus (GEO) database for making the transcriptomic datasets (GSE199939 and GSE80178) publicly available, which laid the foundation for this study.

## DATA AVAILABILITY STATEMENT

The transcriptomic datasets used in this study are publicly available in the Gene Expression Omnibus (GEO) database under the accession numbers GSE199939 and GSE80178. The SASP-related genes (SASPRGs) were obtained from the literature (PMID: 35974106). All other relevant data and analysis code generated in this study are available upon reasonable request from the corresponding author.

## ETHICS STATEMENT

Clinical trial number: not applicable.Ethical approval for this study was granted by the Institutional Review Board of Gansu Provincial Hospital of Traditional Chinese Medicine (approval identifier: GSZY-2025-044-01). All procedures involving human participants were conducted in accordance with the Declaration of Helsinki. Informed consent was obtained from all participants or their legal representatives prior to sample collection.Participants (or legal representatives for incapacitated individuals) signed a consent form acknowledging sample use for research purposes, which was documented and filed by the research team.

---

Table 1. Binding energy of molecular docking between biomarkers and drugs

Table S1. SASP-related genes (SASPRGs)

Table S2. Primer sequence

Table S3. Analysis of differentially expressed genes (DEGs)

Table S4. Gene analysis of key modules

Table S5. Screening analysis of key module genes

Table S6. Candidate genes

Table S7. GO enriched pathways of candidate genes

Table S8. KEGG enriched pathways of candidate genes

Table S9. Pathways enriched by GSEA analysis for CXCR2

Table S10. Pathways enriched by GSEA analysis for SIGLEC7

Table S11. Correlation analysis between biomarkers and differentially infiltrated immune cells

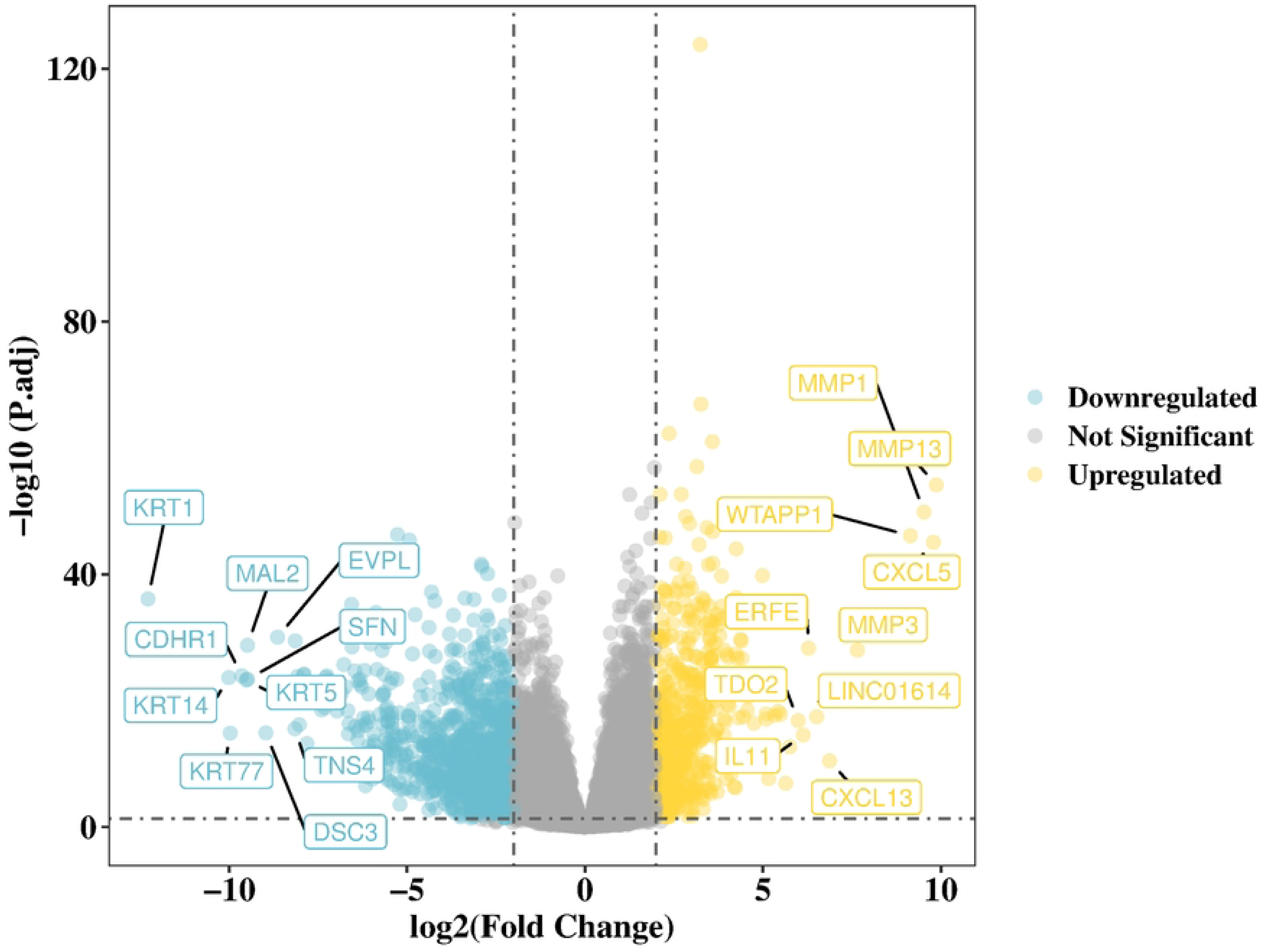

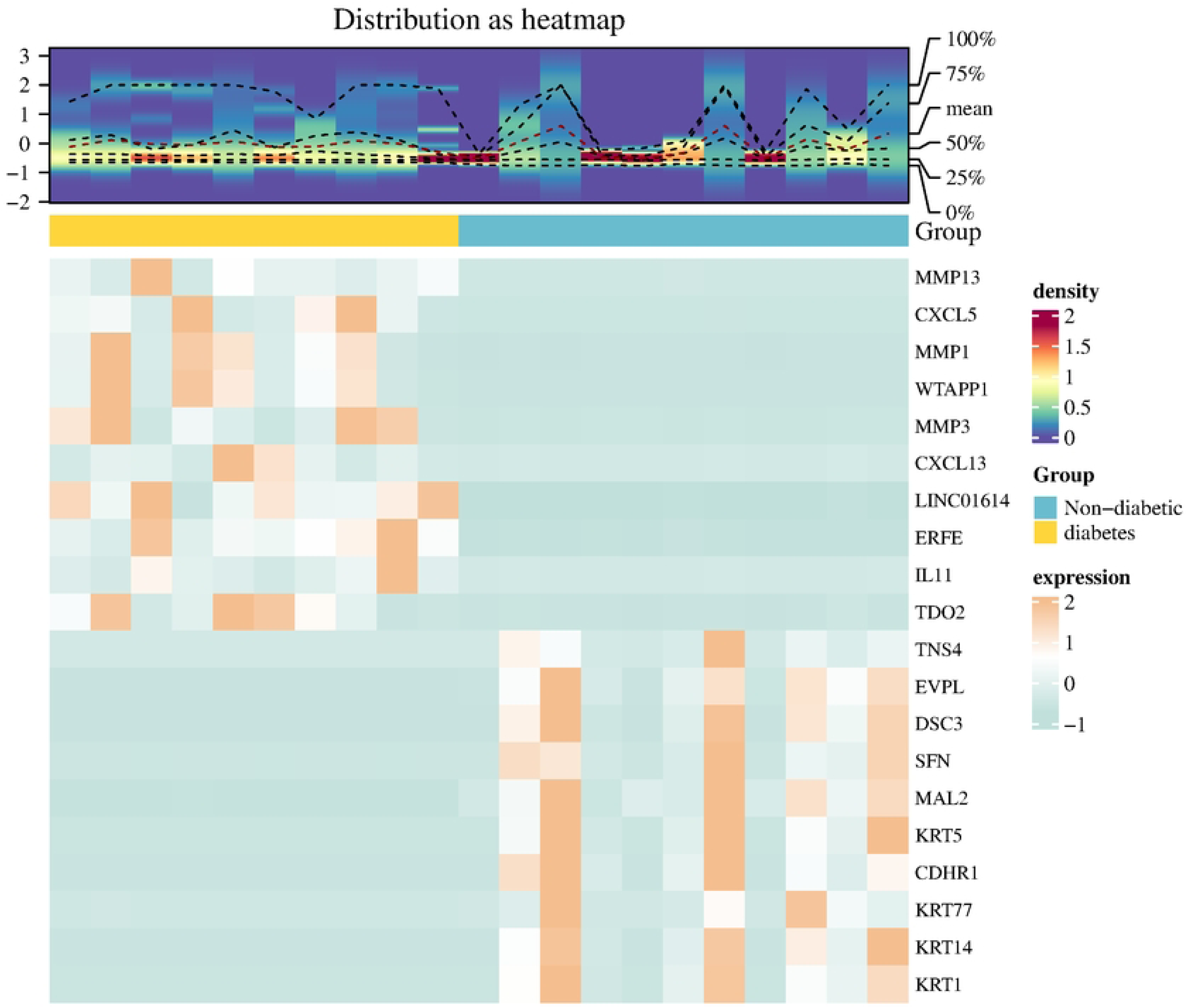

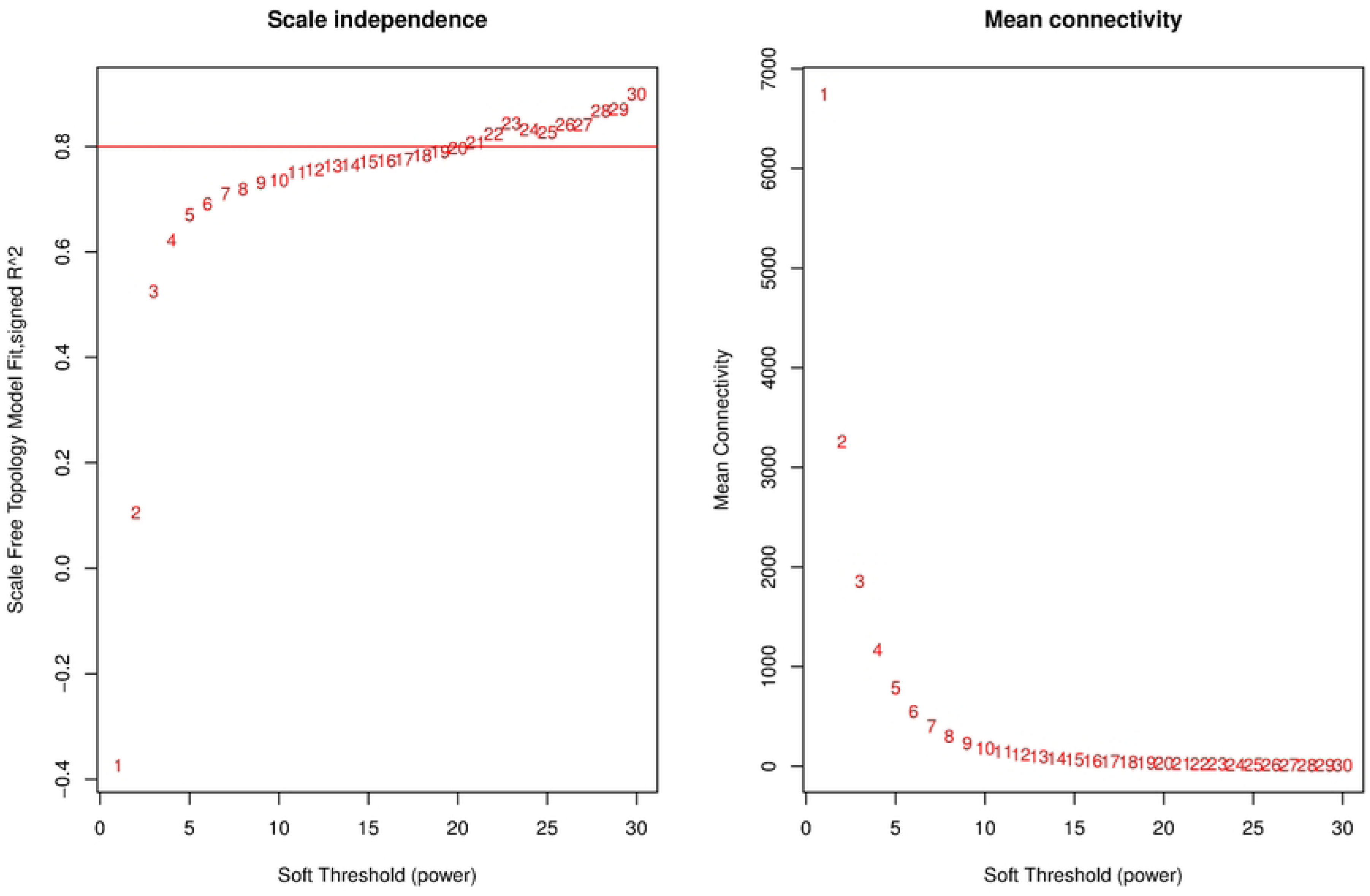

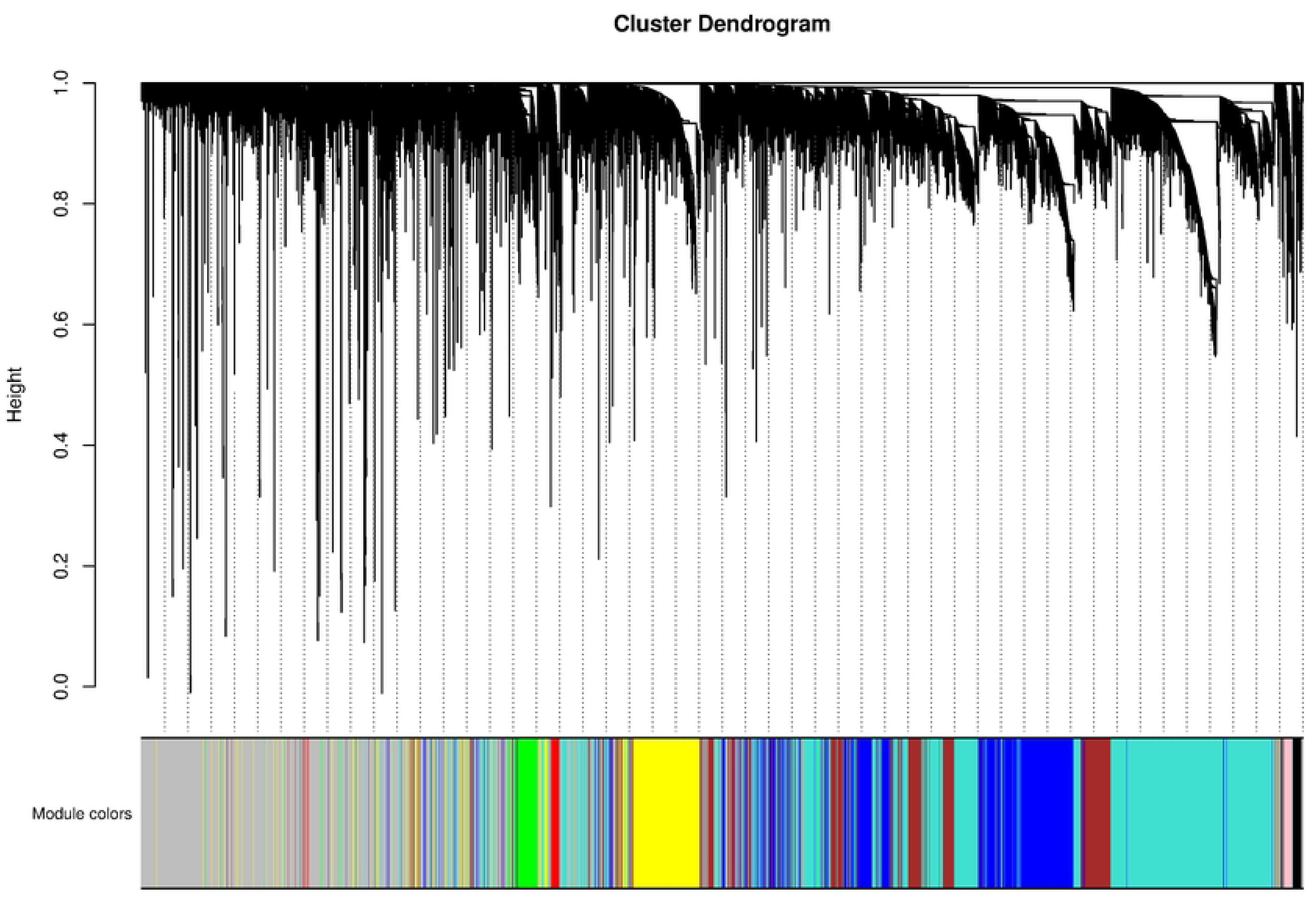

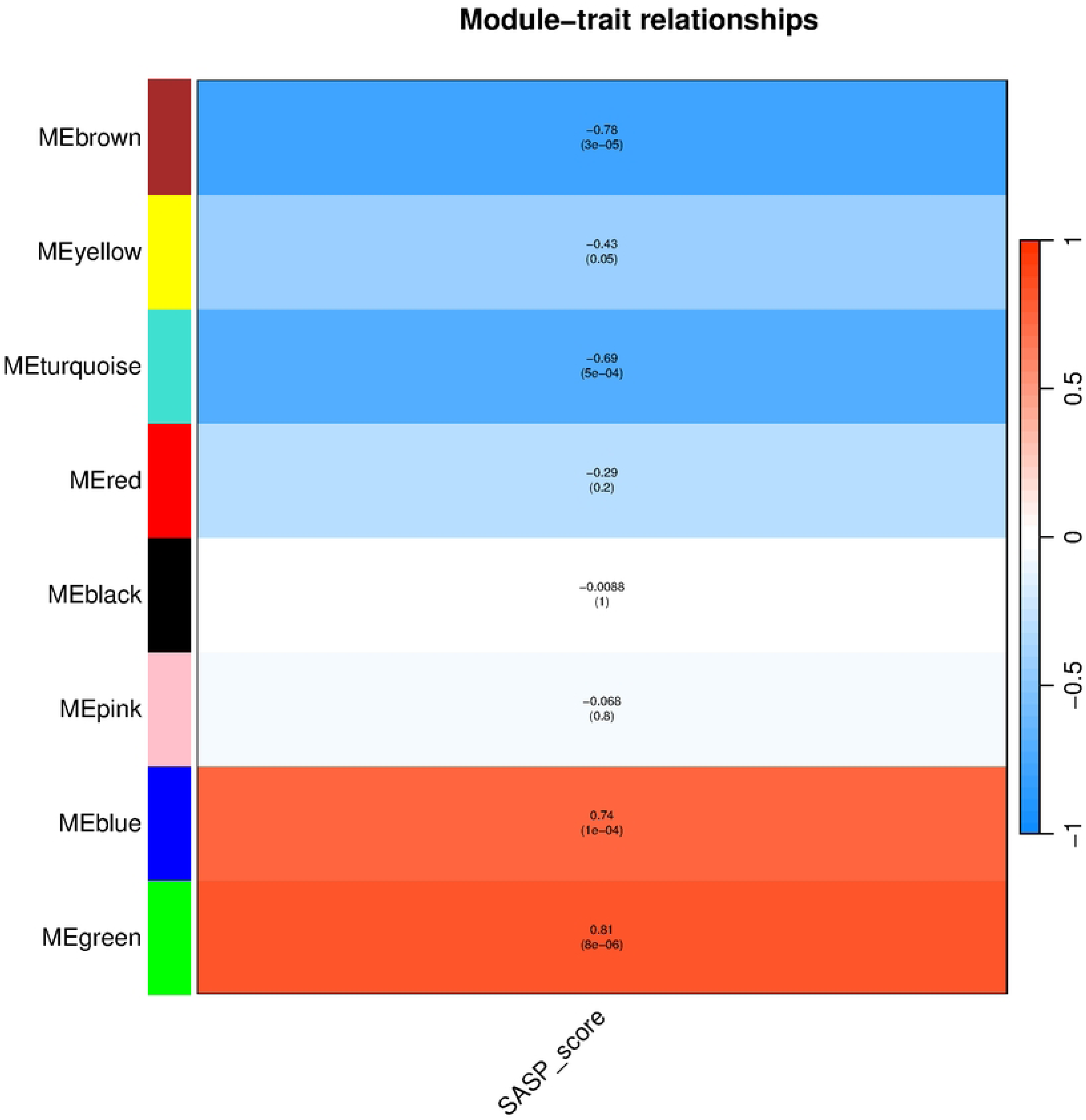

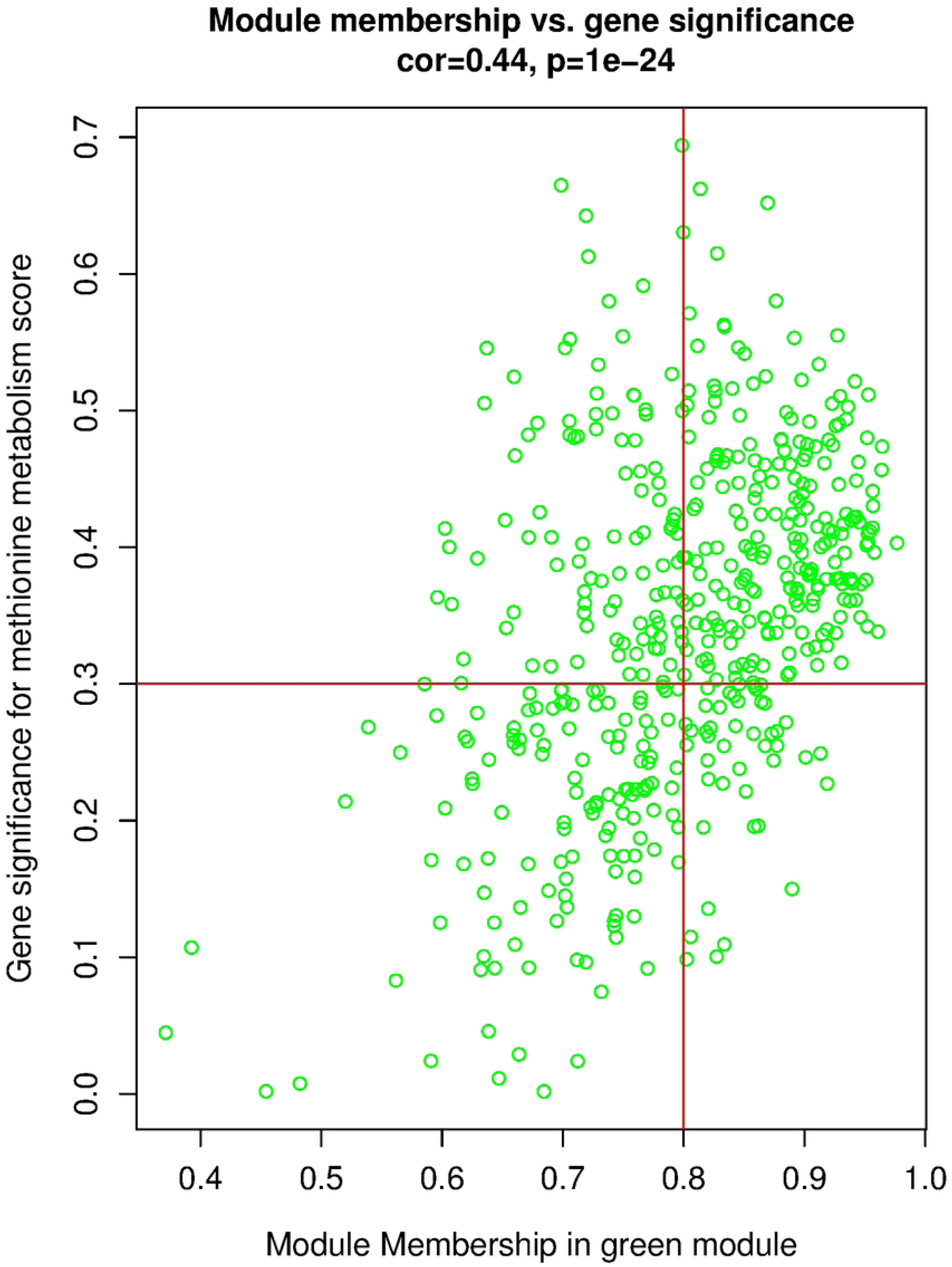

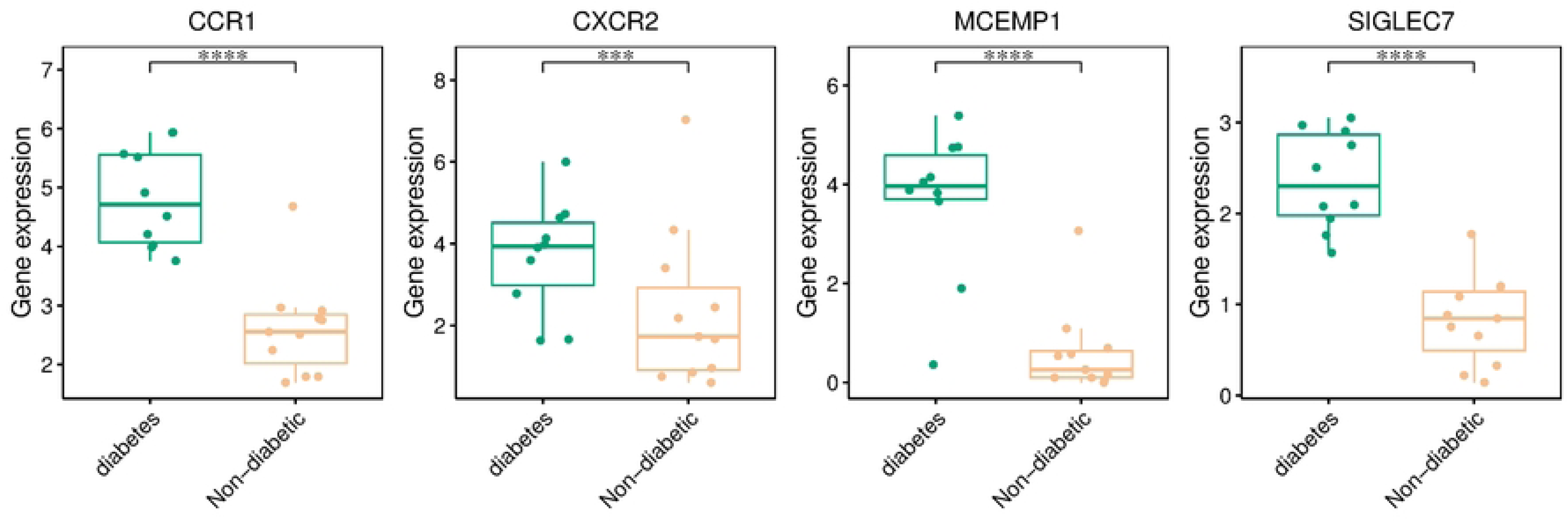

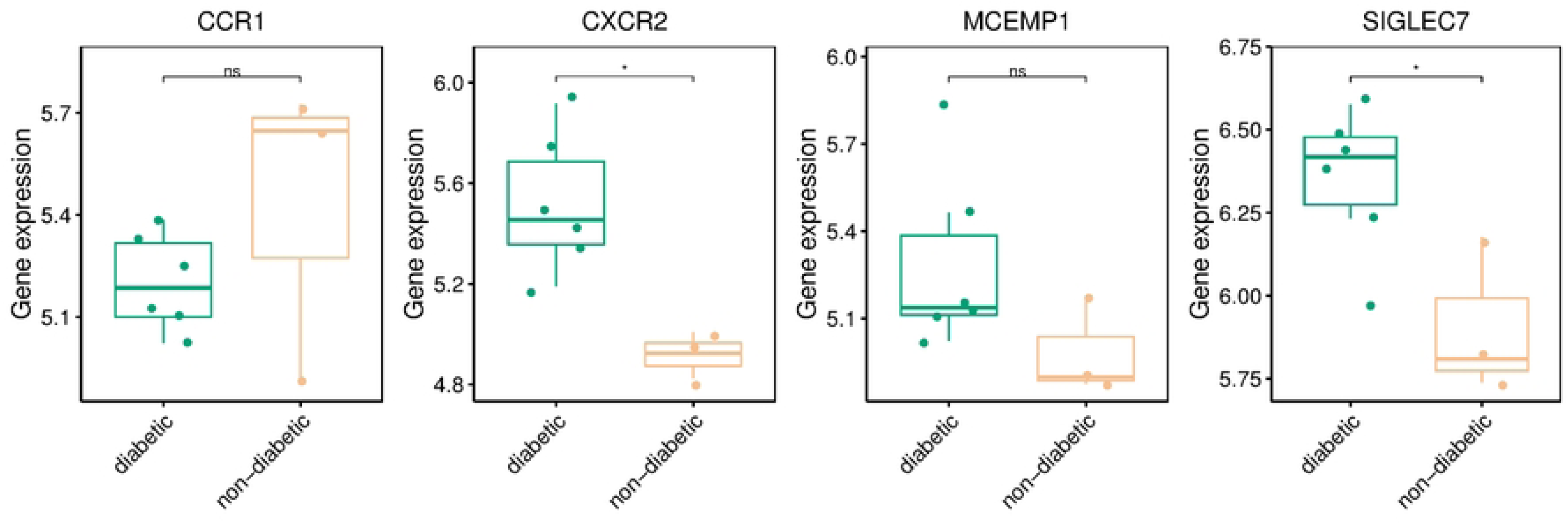

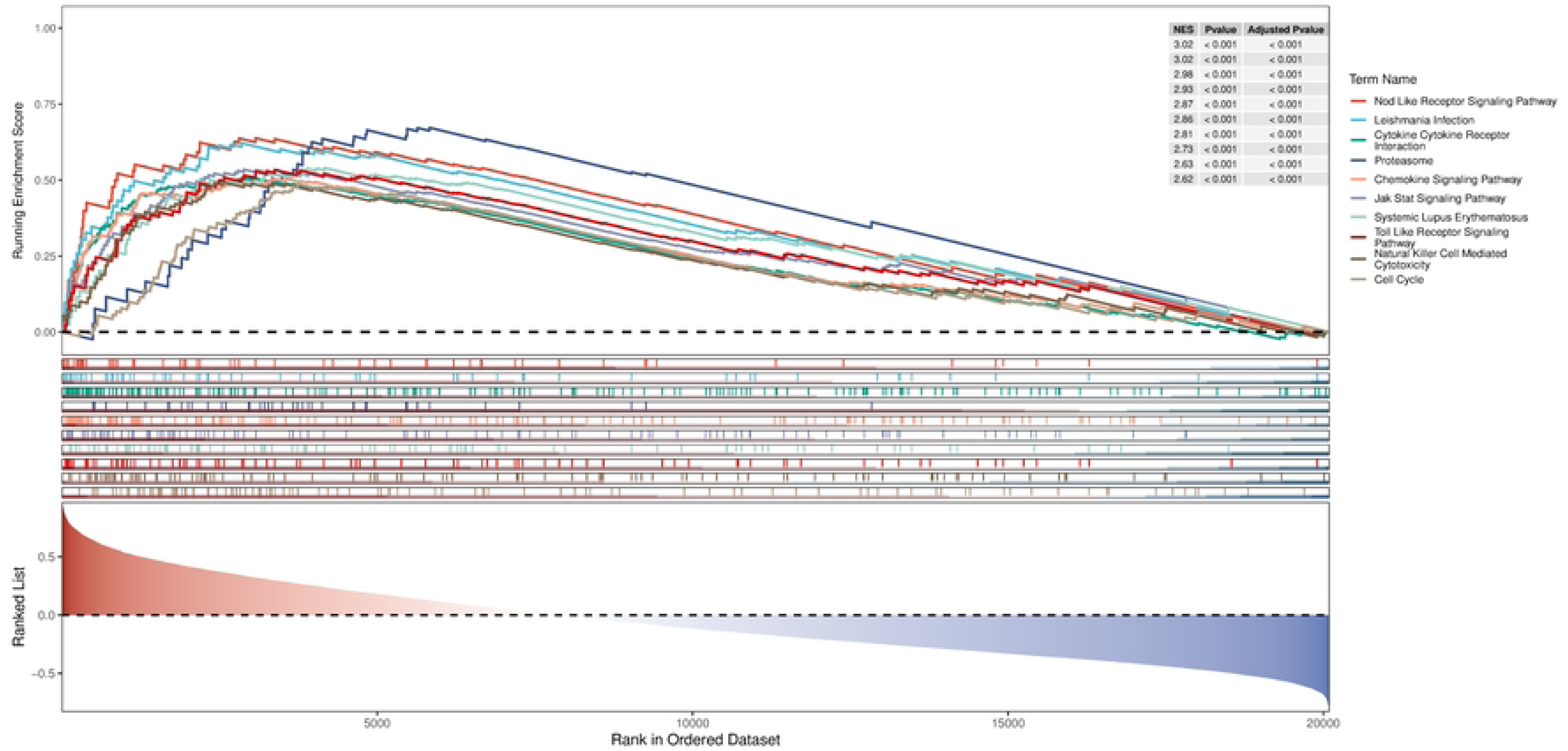

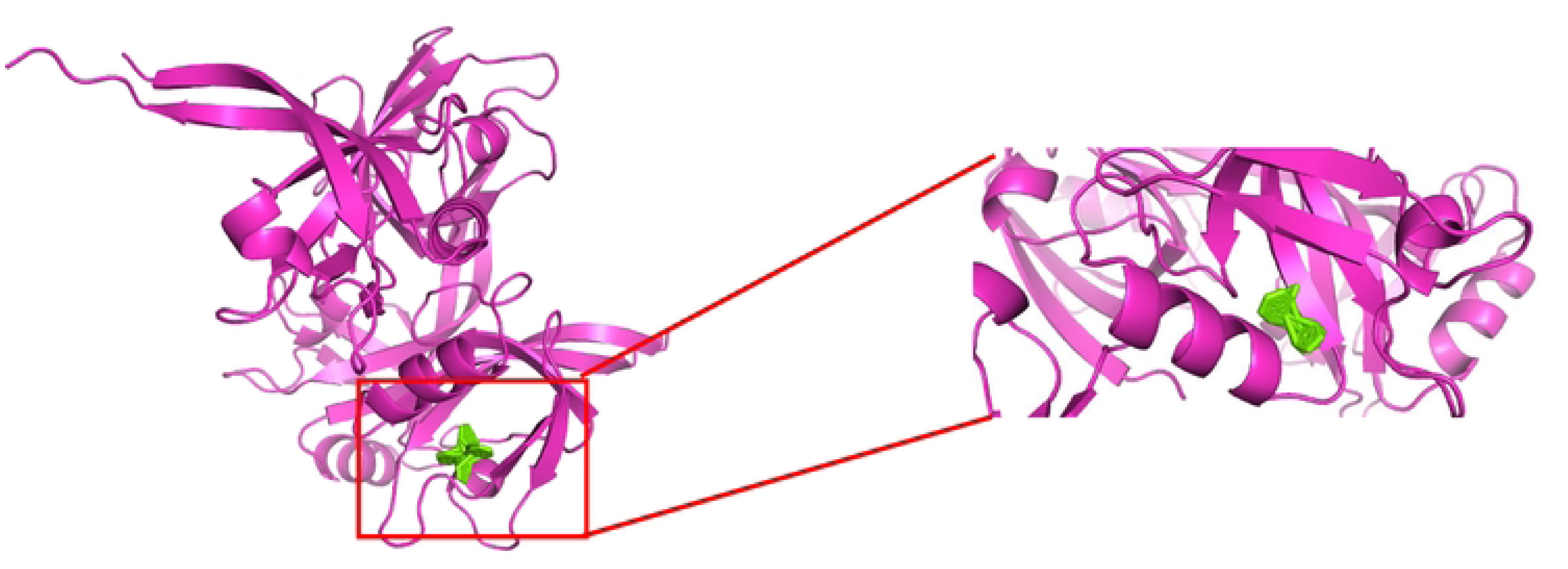

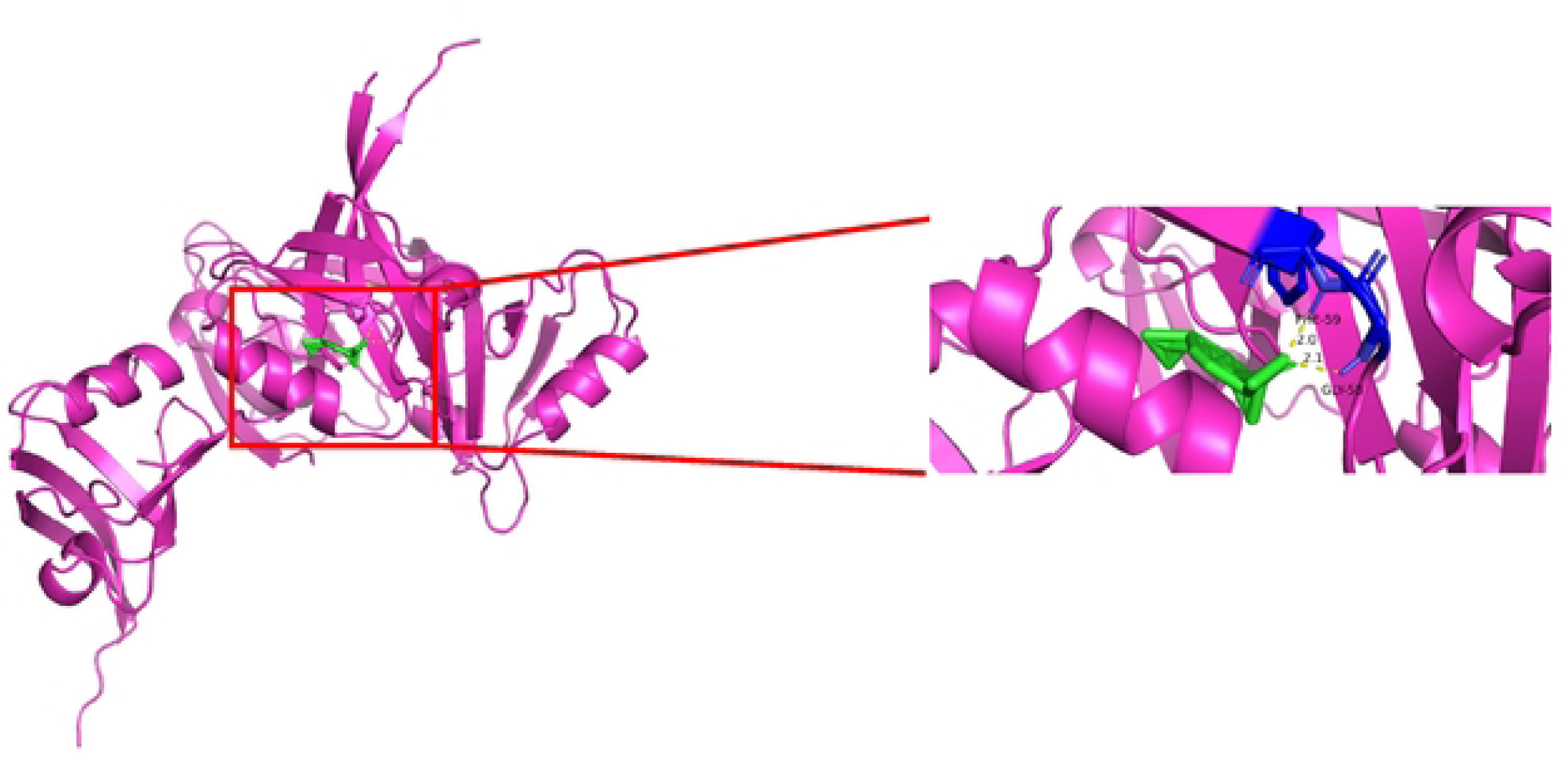

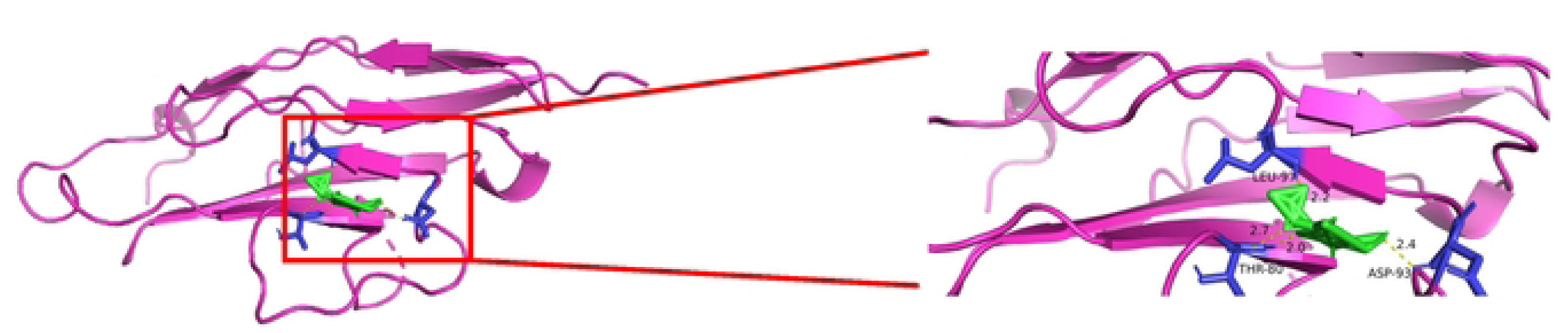

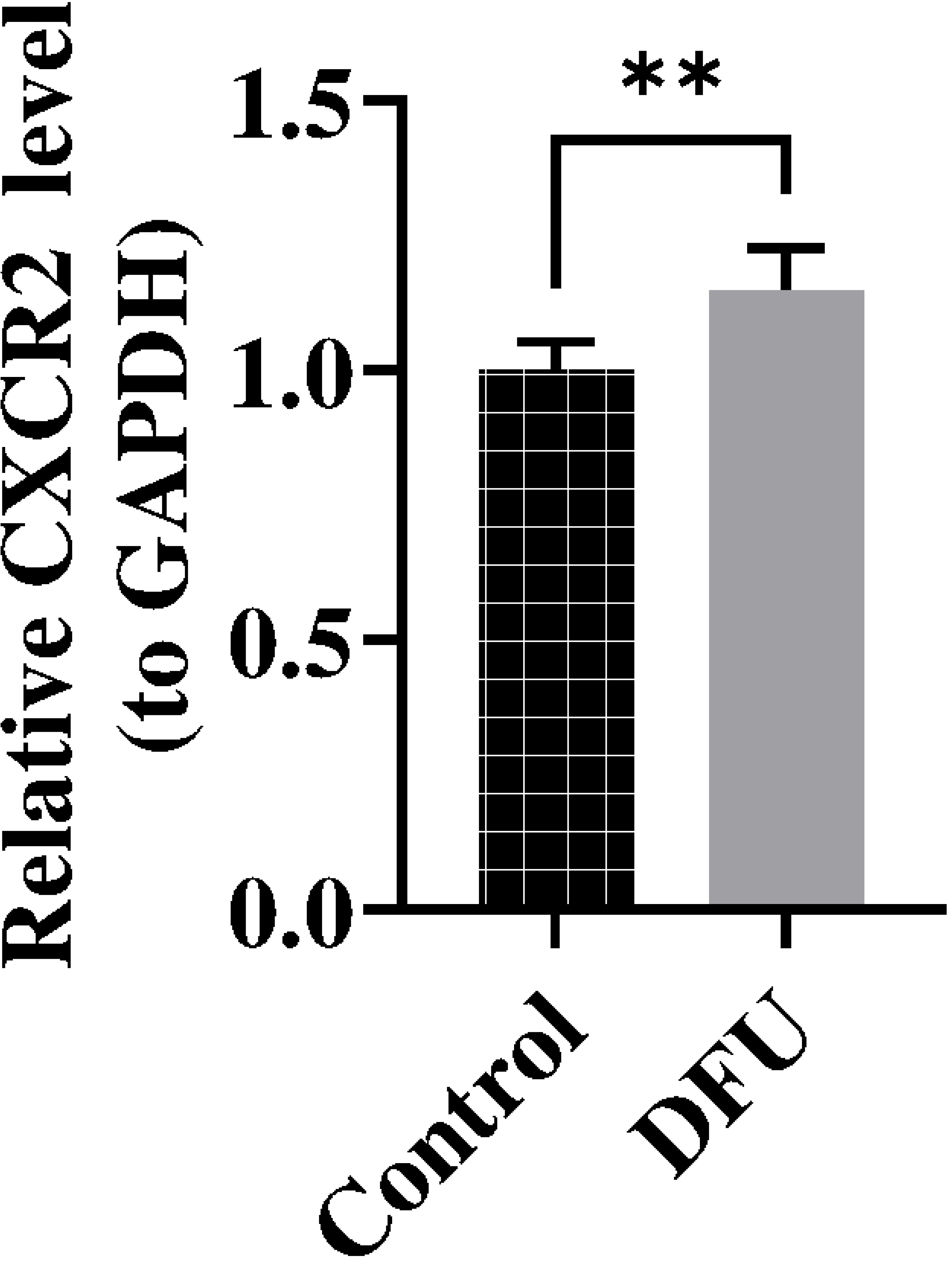

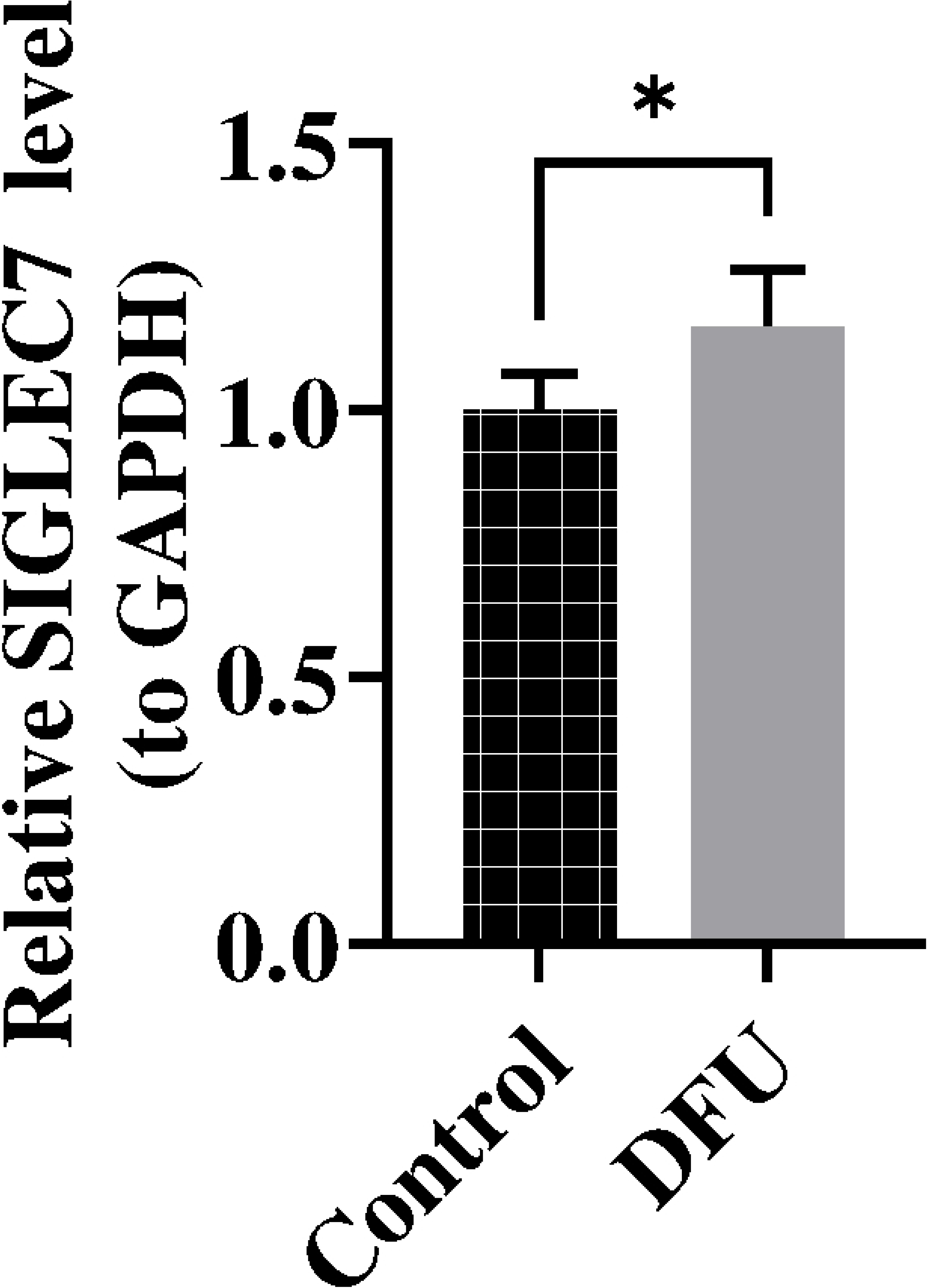

